# S-nitrosylated and non-nitrosylated COX2 has differential expression and distinct subcellular localization in normal and breast cancer tissue

**DOI:** 10.1101/2020.05.20.104612

**Authors:** Sonali Jindal, Nathan D. Pennock, Alex Klug, Jayasri Narasimhan, Andrea Calhoun, Michelle R. Roberts, Rulla M. Tamimi, A. Heather Eliassen, Sheila Weinmann, Virginia F. Borges, Pepper Schedin

**Author notes:** **Corresponding Author:** Pepper Schedin, PhD, Professor Department of Cell, Developmental and Cancer Biology, 2720 SW Moody Ave., Mailing Code: KR-CDCB, Portland, OR, 97201, USA.

## Abstract

Immunohistochemical staining in breast cancer shows both gain and loss of COX2 expression with disease risk and progression. We investigated four common COX2 antibody clones and found high specificity for purified human COX2 for three clones; however, recognition of COX2 in cell lysates was clone dependent. Biochemical characterization revealed two distinct forms of COX2, with SP21 recognizing an S-nitrosylated form and CX229 and CX294 appearing to recognize the same non-nitrosylated COX2 antigen. We found S-nitrosylated and non-nitrosylated COX2 occupy different subcellular locations in normal and breast cancer tissue, implicating distinct synthetic/trafficking pathways and function. Dual stains of ~2000 breast cancer cases show early onset breast cancer has increased expression of both forms of COX2 compared to postmenopausal cases. Our results highlight the strengths of using multiple, highly characterized antibody clones for COX2 immunohistochemical studies and raise the prospect that S-nitrosylation of COX2 may play a role in breast cancer biology.

## Introduction

The cyclooxygenase enzyme COX2, a key mediator of tissue inflammation via prostaglandin production, has been investigated extensively as a cancer biomarker and therapeutic target ^1–4^. Data supporting pro-tumorigenic roles for COX2 include robust preclinical studies identifying COX2 as an oncogene ^5–8^; the demonstration that NSAID-based COX2 blockade inhibits cancer progression in preclinical models ^9,10^; and epidemiologic studies showing that NSAID use correlates with reductions in colon and breast cancer risk ^11–16^. However, prospective clinical trials utilizing aspirin or celecoxib therapy for the prevention, recurrence and treatment of colon ^17–20^ or breast cancer ^14,15,21–23^ show variable results. Further, within the breast cancer field, disparate results of COX2 immunohistochemical (IHC) studies call into question the reliability of COX2 as a breast cancer biomarker or therapeutic target ^24–29^.

In colon, where COX2 is firmly established as a tumor promoter, IHC studies on formalin-fixed paraffin-embedded (FFPE) tissue report minimal COX2 levels in normal epithelium and increased COX2 in at-risk epithelium and invasive cancer ^30–34^. While some breast cancer studies corroborate these results ^35^, others show loss of COX2 in invasive disease compared to adjacent normal tissue ^26,36–39^. One explanation for these divergent results may be methodological, as no standardized approach to COX2 IHC detection has been adopted. For example, one concern regarding αCOX2 antibodies is cross-reactivity, as COX1 is closely related to COX2, with 65% amino acid sequence homology and near-identical catalytic sites ^40,41^.

In this study, we offer a novel explanation for the conflicting data on COX2 expression in breast cancer studies, with implications for other cancers: antibody clone specificity for distinct forms of COX2 based on S-nitrosylation state. First, we investigated the effect of antibody clone on COX1 and COX2 recognition biochemically, focusing on four commonly utilized αCOX2 clones. We validated three clones, SP21, CX229, and CX294 as highly specific for COX2 protein. Unexpectedly, we found these antibody clones differently recognized COX2 in COX2 positive cell lysates, in adjacent normal breast, and in breast and colon cancer tissues. In the breast, we found these distinct staining patterns are due to antibody specificity for S-nitrosylated and non-nitrosylated forms of COX2 respectively. In summary, these studies provide a plausible explanation for disparate COX2 staining patterns observed in breast cancer studies, highlight the strengths of interrogating COX2 with multiple validated antibodies, and demonstrate subcellular localization that infers distinct regulation and function of COX2 based on S-nitrosylation.

## Results

### COX2 Antibody Validations

We assessed four commonly utilized αCOX2 antibody clones, SP21, CX229, CX294, and D5H5 (**Table 1**) for COX2 specificity to recombinant human COX1 and COX2 proteins. We found all four antibodies recognized recombinant human COX2 protein (**Fig 1A, lanes 2-5**). Only D5H5 showed weak reactivity to human COX1 (**Fig 1A, lane 10**) and was eliminated from further evaluation.

**Table 1:** Comparison of structural properties of two COX-2 antibodies

| Designated Clone | SP21 | CX229 | CX294 | D5H5 |
| --- | --- | --- | --- | --- |
| Epitope | Rat COX-2, C-terminus AA sequence 513-604 | Human COX-2, C-terminus AA sequence 580-599 | Human COX-2, C-terminus AA sequence 580-598 | Human COX-2 protein residues surrounding AA sequence 93-123 |
| Host for antibody preparation | Rabbit | Mouse | Mouse | Rabbit |
| Clonality | Monoclonal | Monoclonal | Monoclonal | Monoclonal |
| Epitope Cox-1 overlap | 48AA | 2AA | 2AA | 17AA |
| Vendor Company (catalog #) | Thermo Scientific (RM-9121-R7) | Cayman Chemicals (160112) | Dako (M3617) | Cell Signaling (12282) |

**Figure 1.**
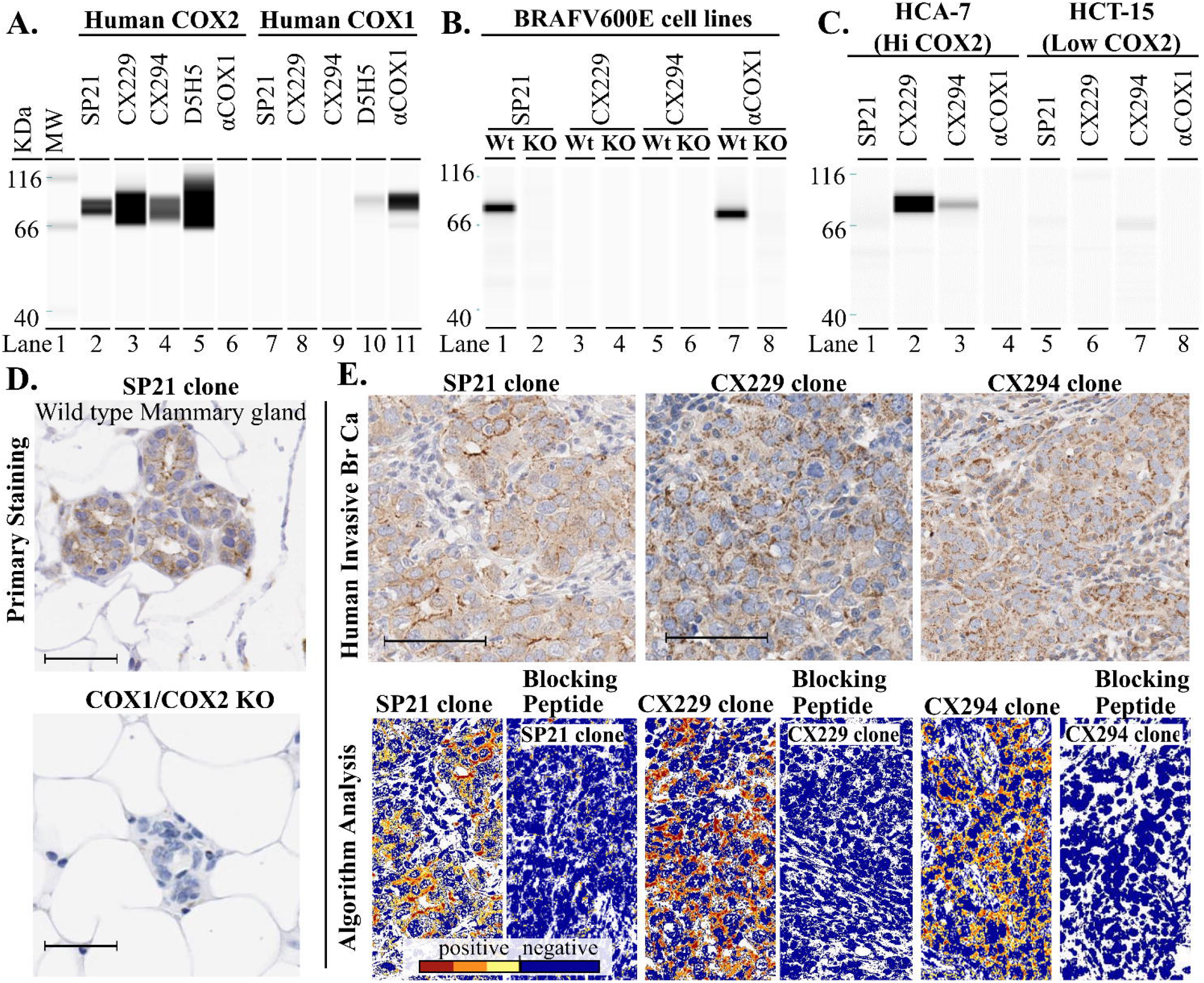
αCOX2 specificity of four distinct antibody clones. **A**. Western blot analysis for αCOX2 clones SP21, CX229, CX294 and D5H5 against recombinant human COX2 and COX1 protein show high specificity of clones SP21, CX229, and CX294 for COX2 protein (A, lanes 2-4, lanes 7-9). Clone D5H5 shows strong reactivity to COX2 protein, but also some reactivity to COX1 protein (A, lanes 5, 10). Lane A11 is the COX1 protein/αCOX1 antibody positive control. **B**. Mouse melanoma BRAFV600E cell lysate is recognized by SP21 in wild type (Wt) cells but not in COX1/COX2 KO cells (B, lanes 1, 2). CX229 and CX294, made against human COX2, do not show reactivity to mouse COX2+ cell lysates (B, lanes 3, 5). **C**. Clones SP21, CX229 and CX294 were probed against human cell lysates with high (HCA-7) and low (HCT-15) COX2 expression. SP21 did not show reactivity to HCA-7 cell lysate (C, lane 1). CX229 and CX294 show reactivity to HCA-7 cell lysate (C, lanes 2, 3). All three clones show minimal to no reactivity to HCT-15 cell lysates (C, lanes 5-7). **D**. FFPE tissue stained using SP21 shows staining in the mammary epithelium of Wt but not in COX1/COX2 KO mice. **E**. Human breast cancer tissue stained for SP21, CX229 and CX294 show robust signal in the tumor cells (E, upper panels, brown stain). Quantitative algorithmic analysis shows positive (orange and red) and negative (yellow and blue) signal for SP21 and CX229 in human breast cancer tissue (E, lower left panels). COX2 epithelial signal with SP21, CX229 and CX294 is blocked using a COX2 specific blocking peptide (E, lower 2^nd^, 4^th^ and 6^th^ panels). Scale bar for all images is 50μm.

We next assessed specificity of SP21, CX229, and CX294 to detect COX2 protein in cell lysates from mouse melanoma BRAFV600E cells with wild type or genetically deleted COX1/COX2 (KO) ^42^. SP21 detected COX2 protein in wild type but not in KO cells (**Fig 1B, lanes 1-2**). CX229 and CX294 did not detect murine COX2 (**Fig 1B, lanes 3 and 5**), consistent with reported human specificity for these clones. To assess antibody reactivity to human COX2 protein, we utilized human colon cancer cell lines with high (HCA-7) and low (HCT-15) COX2 expression ^43,44^. As anticipated, CX229 and CX294 detected COX2 protein in the high COX2 expressing HCA-7 cells (**Fig 1C, lanes 2-3**), but not in the low expressing HCT-15 cells (**Fig 1C, lanes 6-7**). Unexpectedly, SP21 did not detect COX2 protein in the high COX2 expressing HCA-7 cell lysate (**Fig 1C, lane 1**), even though SP21 robustly detects human recombinant COX2 protein (**Fig 1A, lane 2**) ^27,45^.

We next assessed for COX2 antibody specificity in FFPE tissues. Because SP21 recognizes murine COX2, we utilized genetically modified mouse models to confirm antibody specificity ^38^. We found that SP21 recognized murine COX2 in mouse mammary epithelial cells in wild type but not in COX2 KO glands, confirming specificity (**Fig 1D**). Next, a previously validated COX2 positive human breast cancer case ^38^ was selected to assess SP21, CX229 and CX294 staining. All three clones stained this COX2 positive control tissue (**Fig 1E**). We next demonstrated SP21, CX229 and CX294 specificity for COX2 by confirming that the majority of antibody signal was lost with the addition of a COX2 specific blocking peptide (**Fig 1E**). Thus, all three COX2 clones show high specificity and sensitivity for COX2 in FFPE tissues.

### S-nitrosylated and non-nitrosylated forms of COX2

Given the high specificity and sensitivity of SP21, CX229 and CX294 for COX2 protein, it is unclear why SP21 would not recognize COX2 in the COX2 high expressing HCA-7 cells (**Fig 1C, lane 1**). To address this question, we examined the amino acid sequences used as immunogens (**Fig 2A**) for SP21, CX229 and CX294 antibody generation. We found a post-translation modification site for S-nitrosylation at Cys-526 only in the SP21 immunogen (**Fig 2A, red arrow**). CX229 and CX294 were made using essentially identical amino acid sequences. Since they similarly recognized COX2 in human colorectal cancer cell lines by western blot and have essentially identical staining in a breast cancer TMAs (n = 56, **Supplemental Fig 1 and Supplemental Table 1**), we focused our subsequent biochemical analyses on SP21 and CX229. To address whether SP21 preferentially recognizes S-nitrosylated COX2, we employed the strategy of biochemically adding and removing nitric oxide moieties to COX2 protein and then assessing antibody recognition by western blot. To obtain a source of COX2 that is differentially recognized by SP21 and CX229 and suitable for S-nitrosylation modifications, we performed COX2 immunoprecipitation of HCA-7 cell lysates using CX229, as CX229 recognizes COX2 in this cell line (as does CX294) whereas SP21 does not (**Fig 1C, lane 2 vs. lane 1**). As expected, CX229 recognizes the COX2 protein immunoprecipitated by the CX229 antibody (**Fig 2B, lane 1**), while CX229-immunoprecipitated COX2 was undetected by SP21 **(Fig 2B, lane 2**). Since the SP21 immunogen includes the putative COX2 S-nitrosylation site, we reasoned that HCA-7 COX2 protein is non-nitrosylated, and that S-nitrosylation might convert HCA-7 COX2 to an SP21-recognizable form. To test this idea, the above CX229 immunoprecipitated COX2 was S-nitrosylated by incubation with S-nitrosoglutathione (SNOG) ^46^. We found that SP21 detected HCA-7 COX2 only after incubation with S-nitrosoglutathione (**Fig 2B, lane 4**), which is consistent with SP21 specifically recognizing an S-nitrosylated form of COX2.

**Figure 2.**
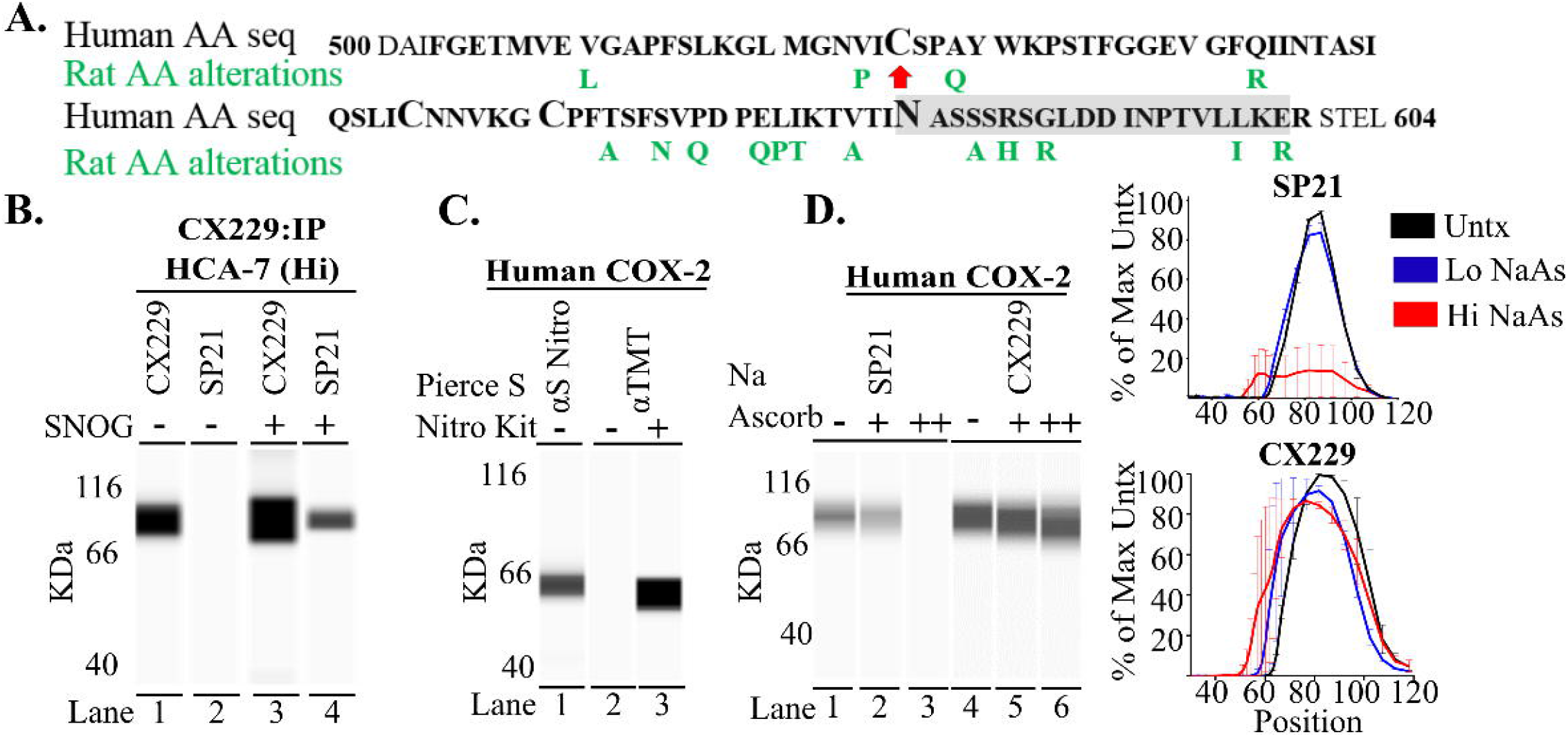
SP21 recognizes S-nitrosylated COX2. **A.** Human and Rat amino acid (AA) sequence of PTGS2 gene region used as immunogens (SP21= black bold, CX229= gray box). Potential post-translational modification (larger font size) S-nitrosylation site is seen at cysteine 526 (red arrow), disulfide bond sites at AA 555 and 561, and a glycosylation site at AA 580. **B**. COX2 immunoprecipitation (IP) of HCA-7 cell lysate using CX229 is recognized by CX229 (B, lane 1) but not SP21 (B, lane 2). On biochemical S-nitrosylation of the CX229 IP, SP21 regains reactivity to COX2 (B, lane 4). **C**. Western blot analysis confirms recombinant human COX2 protein contains the S-nitrosylated form of COX2 as detected by a pan-nitrosylation specific monoclonal antibody (C, lane 1) as well as by assessment of nitrosylation modification using the Pierce S-nitrosylation kit and the anti-TMT antibody (C, lanes 2-3). **D**. Western blot analysis confirms dose-escalating Na ascorbate treatment (−, +, ++) used for de-nitrosylation of recombinant human COX2 protein results in dose dependent loss of SP21 signal (D, lanes 1-3). No or minimal loss of CX229 reactivity was observed after Na ascorbate treatment (D, lanes 4-6). Quantitation of COX2 western blots from three separate Na ascorbate experiments (D, right panel electrophoretograms).

To determine if SP21 antibody signal is lost with de-nitrosylation of COX2 protein, as predicted if SP21 is specific for S-nitrosylated COX2, we next performed de-nitrosylation assays. Because recombinant human COX2 is recognized by SP21, suggesting S-nitrosylation (**Fig 1A, lane 2**), we first determined if the recombinant human COX2 is S-nitrosylated using a pan-nitrosylation specific monoclonal antibody (**Fig 2C, lane 1**), and a commercial (Sigma) biochemical detection kit for S-nitrosylation (**Fig 2C, lane 2&3**). Both methods demonstrate that purified recombinant human COX2 contains S-nitrosylated COX2. We then utilized Na ascorbate to de-nitrosylate recombinant human COX2, utilizing a methodology previously reported in mouse cell lysates ^47^. Na ascorbate treatment resulted in a dramatic dose-dependent decrease in SP21 signal (**Fig 2D, lane 3, & upper electrophoretogram**). In contrast, de-nitrosylation of recombinant COX2 did not significantly reduce the CX229 signal (**Fig 2D, lane 6, & lower electrophoretogram**). These western blot assay data are consistent with SP21 specifically recognizing S-nitrosylated COX2, whereas CX229 signal appears independent of S-nitrosylation.

### S-nitrosylated and non-nitrosylated COX2 staining patterns in human breast and colon cancer tissue

We next sought to determine if the S-nitrosylation state of COX2, as detected by SP21 and CX229, could result in disparate staining results in cancer tissue. To this end we stained sections of human breast and colon TMAs, with 52 and 53 cores respectively, using routine chromogen-based IHC methods. In breast TMAs, 2% of the cores stained preferentially with SP21, whereas 35% of the cores stained preferentially with CX229, with little to no overlap between staining patterns (**Fig 3A & Supplemental Table 2**). Further, colon TMAs also had differential staining patterns for SP21 and CX229, suggesting relevance beyond breast (**Fig 3A & Supplemental Table 2**). While unique stating patterns for SP21 were anticipated based on its specificity for S-nitrosylated COX2, the fact that CX229 antibody signal did not overlap with the SP21 signal was unanticipated and strongly suggests that SP21 and CX229 recognize distinct forms of COX2.

**Figure 3.**
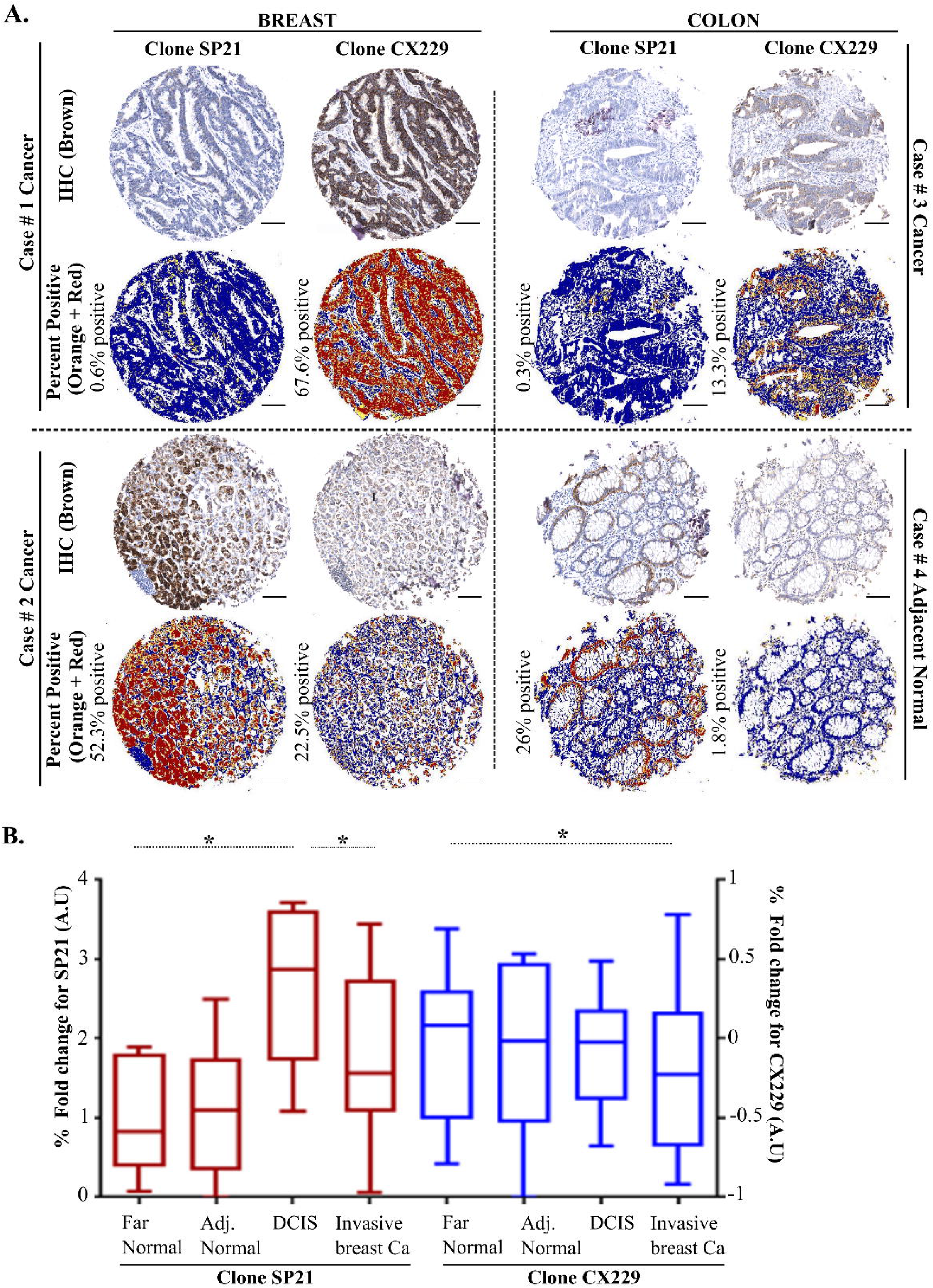
Two αCOX2 clones show differential pattern of staining. **A.** TMA cores from breast and colon cancer cases show preferential CX229 (cases #1 & #3) or preferential SP21 staining (cases #2 & #4). Algorithmic analysis for each TMA core shows COX2 positive signal (orange and red) compared to negatively stained tissue (blue). **B**. Sequential sections of individual cases were stained for SP21 (red) and CX229 (blue) and evaluated for COX2 expression in far and adjacent normal, DCIS, and Invasive breast cancer tissue (n=9). SP21 shows highest expression in the DCIS lesions. Clone CX229 shows highest COX2 expression in normal far and adjacent breast epithelium, with decreased COX2 expression in invasive cancer (P values: *≤0.05). Scale bar for all images is 100μm.

### SP21 and CX229 show opposing staining trends in normal adjacent, DCIS and invasive breast cancer tissue

We next addressed antibody staining patterns in breast tissues that contained far normal, adjacent normal, DCIS, and invasive tumor on a single slide. The SP21 staining pattern was consistent with the paradigm of increased COX2 expression in DCIS compared to adjacent normal tissue (**Fig 3B, Supplemental Fig 2A**). In this small cohort, we also saw a significant decrease in SP21 COX2 signal in invasive breast cancer compared to adjacent DCIS lesions; however, SP21 staining in invasive breast cancer trended higher than in adjacent normal tissue. In contrast, with CX229, the highest COX2 expression was observed in histologically normal epithelium, with modest but progressive loss of COX2 expression in invasive cancer (**Fig 3B, Supplemental Fig 2B)**. These data provide preliminary evidence that the S-nitrosylated and non-nitrosylated forms of COX2 are both present in breast cancer cases, with differentially elevated expression during breast cancer progression. Importantly, these data show how clone selection for COX2 antibody may yield substantially different results in breast cancer studies and demonstrate the inherent limitations of assessing COX2 expression using a single antibody clone.

### Distinct intracellular localization of S-nitrosylated COX2

With the biochemical confirmation that SP21 and CX229 differentially recognize COX2 protein based on S-nitrosylation state, we next assessed where these two forms of COX2 localize within human breast cancers. To obtain information about COX2 localization at a sub-cellular resolution, we performed dual immunofluorescence (IF) staining with SP21 and CX229, as well as with SP21 and CX294. We found that individual cancer cases could be dominated by SP21 signal (**Fig 4A, left panel**), CX229 signal (**Fig 4A, middle panel**) or both (data not shown). Additionally, SP21 and CX294 dual stained cases looked nearly identical to cases stained with SP21 and CX229 (Fig 4 and **Supplemental Fig 4**). Of note, in tumors and adjacent normal breast tissue there was virtually no overlap in cellular localization of SP21 and CX229 nor SP21 and CX294 signal, providing further evidence that SP21 and CX229/CX294 recognize distinct forms of COX2 (**Fig 4A, Supplemental Fig 4**). Further in adjacent normal tissue, SP21 and CX229/CX294 stained distinct subcellular regions. Specifically, within acinar structures CX229 and CX294 dominantly stained lateral plasma membranes (**Fig 4A, right panel green signal, Supplemental Fig 4**), whereas SP21 primarily stained apical junctional regions (**Fig 4A, right panel red signal, Supplemental Fig 4**). These data are consistent with distinct trafficking and function of S-nitrosylated and non-nitrosylated COX2 in adjacent normal tissue.

**Figure 4.**
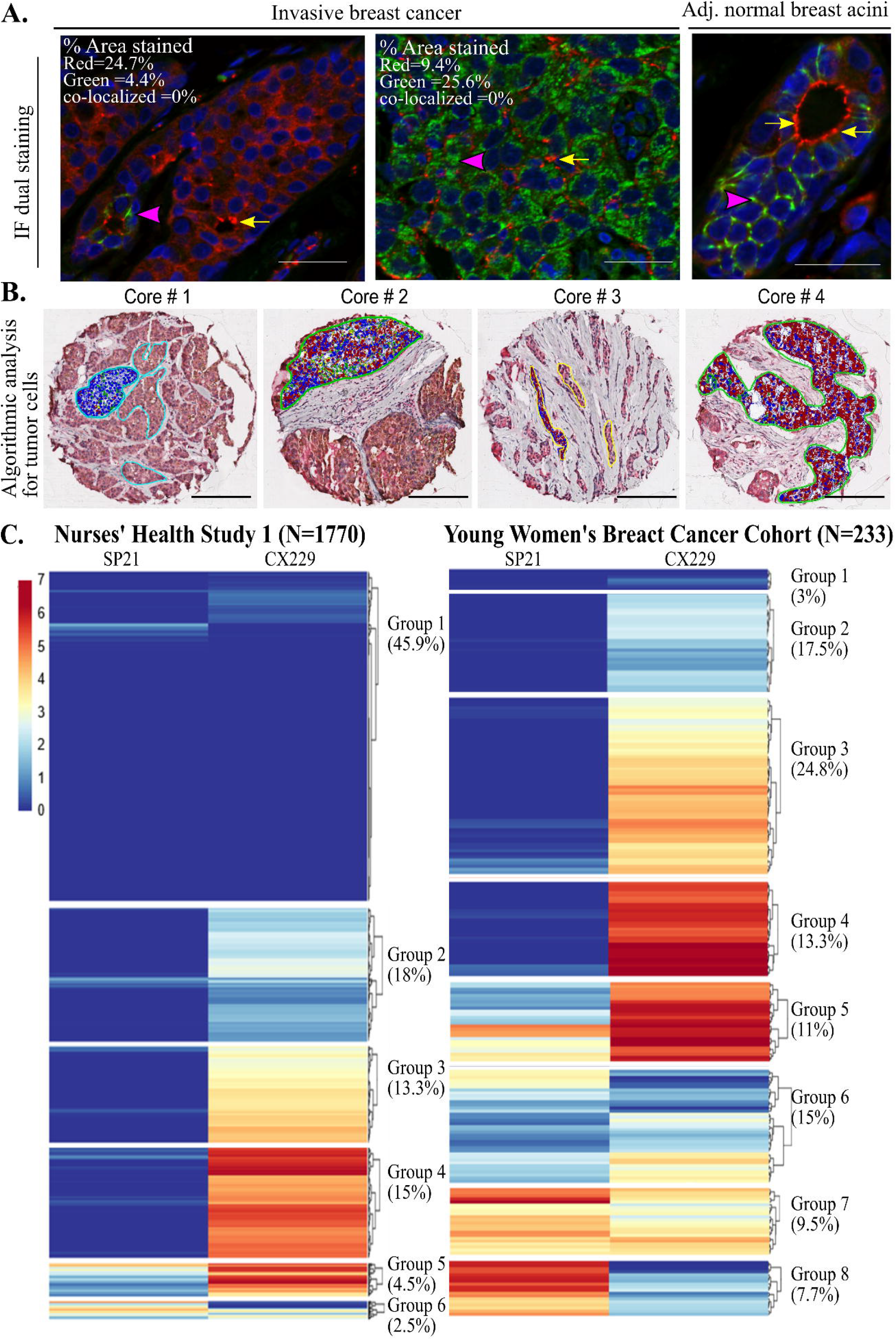
Variation of SP21 and CX229 expression in two breast cancer cohorts. **A.** IF staining of two invasive cancer cases show predominant staining for either SP21 (left panel) or CX229 (middle panel) and absence of co-localization (SP21 = red, CX229 = green, co-localization = yellow). Percent stained area for each clone and percent co-localization are listed within the images. Similarly, IF staining of normal breast acini show SP21 (red signal) and CX229 (green signal) stain distinct cellular locations with minimal overlap (yellow signal, right panel). Pink arrow heads show intense localized staining for CX229. Yellow arrows show intense localized staining for SP21. Scale bar =20 μm **B.** TMA cores with dual staining for SP21 and CX229 were dual stained and annotated for tumor epithelium. The algorithmic analyses of positive staining for SP21 (red), CX229 (green), colocalization (yellow) and negative (blue) is shown. Scale bar is 100μm **C**. Hierarchical clustering analysis for SP21 and CX229, assessed independently for each breast cancer cohort using R studio, shows cohorts separate into distinct groups. NHS1 breast tumor cores (N=8612, 1770 cases) clustered into 6 COX2 expression groups with the largest group (n=3961, 45.9%) exhibiting very low expression for both SP21 and CX229. Groups 2 (n=1580, 18%), 3 (n=1154, 13.3%), and 4 (n=1308, 15%), were defined by cores with low, medium and high expression of CX229 respectively, and very low SP21 expression. Cluster 5 (n=389, 4.5%) had medium level of SP21 expression and high CX229 expression and cluster 6 (n=220, 2.5%) contained cores with medium expression of SP21, but low expression of CX229 (left panel). The young women breast cancer study clustered into 8 groups with cluster 1 (n=7, 1.8%) with very low expression of both SP21 and CX229. Groups 2 (n=34, 17.5%), 3 (n=58, 24.8%) and 4 (n=31, 13.3%) had very low SP21 and low, medium and high CX229 expression respectively. Groups 5 (n=26, 11%) and 6 (n=37, 15%) had high and low CX229 expression respectively with low to medium SP21 expression. Groups 7 (n=22, 9.5%) and 8 (n=18, 7.7%) had medium to high SP21 and low CX229 expression (right panel).

### Variation of SP21 and CX229 Staining in Two Large Breast Cancer Cohorts

To further understand the inter-case variation between S-nitrosylated and non-nitrosylated COX2 expression in breast cancer, we stained two large breast cancer cohorts, the Nurses Health Study 1 (NHS1) cohort and the University of Colorado Young Women’s Breast Cancer Translational Program (YWBCTP) cohort, with ~2000 combined cases (**Supplemental Table 3**). We used an optimized dual SP21 and CX229 staining protocol where we confirmed that neither antibody order nor chromogen selection significantly impacted staining results (**Supplemental Fig 3**). We found ~95% of the COX2 signal comes from tumor cells compared to stromal cells, results consistent with previous reports ^38^. To account for differences in stromal composition between cases, we restricted analyses to tumor cells by positively annotating tumor cell clusters followed by computer-assisted quantitation of antibody signal (Aperio ImageScope analysis software (Leica Biosystems, Vista, CA) (**Fig 4B**)). To this end, SP21 and CX229 COX2 expression for each sample was compiled as a continuous variable and hierarchical clustering was performed using R studio software. Independent K-mean clustering for SP21 and CX229 in both cohorts showed nearly identical cutoff values for positivity (SP21: NHS-2.9%, YWBCTP-2.8%, CX229: NHS-6.1%, YWBCTP-6.2%) indicating similar staining intensity, which permits comparisons between the two cohorts.

The NHS1 cohort is primarily composed of women diagnosed with breast cancer later in life, with an average age at diagnosis of 58 years, and consists of N = 8612 cores representing 1770 cases. By hierarchical clustering, the NHS1 breast tumor cores clustered into 6 COX2 expression groups with group 1 (45.9% of cores), the largest group, exhibiting very low expression for both SP21 and CX229 (**Fig 4C, group 1**). Group 2 (18% of cores), group 3 (13.3% of cores), and group 4 (15% of cores), were defined by cores with low, medium and high expression of CX229 respectively, but very low SP21 expression. Group 5 (4.5% of cores) is defined by medium levels of SP21 expression and high CX229 expression. Finally, group 6 (2.5% of cores) contained cores with medium expression of SP21, but low expression of CX229. Since COX2 expression in the normal breast has been demonstrated to be hormone dependent ^38^, we next assessed the expression of SP21 and CX229 in a cohort of young women’s breast cancer (YWBCTP, N = 233 cases) with an average age at diagnosis of 38 years. In the YWBCTP cohort, only 3% of cases had very low expression of CX229 and SP21 (**Fig 4D, group 1**) compared to 45.9% of cores in the NHS1 cohort (**Fig 4C, group 1**). In particular, SP21 was much higher in the YWBCTP cohort, which resulted in the formation of two additional groups with very high expression of SP21 in ~17% of cases (**Fig 4D, groups 7 and 8**). In sum, COX2 expression, for both S-nitrosylated and non-nitrosylated COX2 varies widely between patients, with non-nitrosylated COX2 being more commonly expressed than the S-nitrosylated form. Additionally, expression of both S-nitrosylated and non-nitrosylated COX2 is more common in the YWBCTP cohort of early onset breast cancers compared to the older, NHS1 cohort.

## Discussion

In this study, we identify commonly used COX2 antibodies that differentially recognize distinct forms of COX2 based, at least in part, on post-translational S-nitrosylation. Evidence of distinct cellular synthetic, trafficking, and functional pathways for these two forms of COX2 is suggested by non-overlapping staining patterns in adjacent normal breast and cancer tissues. Similar staining patterns were observed in colon cancer, which supports relevance of these findings in colon cancer. Further, since COX2 biology is thought to be important in many cancer types and diseases, our discovery of S-nitrosylation state specific COX2 antibody clones is likely to have broad impact. Additionally, in a small cohort, we observed the S-nitrosylated form of COX2 is highest in DCIS lesions compared to adjacent normal breast tissue, with levels decreased in invasive breast cancer. This pattern of expression is similar to that observed for HER2, a bona fide breast cancer oncogene that is expressed at highest levels in DCIS lesions ^48^. Overall, high expression of S-nitrosylated COX2 in DCIS lesions is an observation consistent with the established paradigm of COX2 as pro-tumorigenic. In contrast, levels of non-nitrosylated COX2 were unchanged between adjacent normal and DCIS, with a modest decrease in invasive tumor. These disparate staining patterns, based on S-nitrosylation state of COX2, may help explain why previous breast cancer studies report that COX2 expression either directly ^4,26,49,50^, inversely ^38^ or failed to correlate with breast cancer risk, progression and or outcomes ^25,27^. While we are the first to validate antibody reagents that distinguish COX2 based on its S-nitrosylation state, the impact of COX2 S-nitrosylation in the context of breast cancer remains to be determined.

COX2 S-nitrosylation is dependent on nitric oxide (NO) biology, and data from the NO field provides strong rational for pursuing a potential role for COX2 S-nitrosylation in breast cancer. In endothelial and neuronal cells, NO is produced via expression of the NO synthases eNOS and nNOS, which regulate physiologic vasodilation and neuronal signaling, respectively ^51^. Evidence for an inducible form of NO synthase (iNOS) was first reported in macrophages ^52^, and led to the discovery of NO as a principal regulator of tissue inflammation ^51^ with likely roles in cancer ^51,53^. For example, in triple negative breast cancer (TNBC) cell lines, defined as ER, PR and Her-2 negative, iNOS signaling promotes stem-like properties and metastatic potential ^54^. Further, iNOS blockade as a single agent reduces TNBC growth, metastasis ^55^ and enhances efficacy of chemotherapy in xenograft models ^56^. Importantly, these pre-clinical studies define a novel, NO-centric path toward the possible treatment of aggressive TNBC. Interrogating COX2 S-nitrosylation by breast cancer subtype, grade, and stage is a potentially fruitful next step for understanding COX2 as a biomarker of risk as well as therapeutic target.

Evidence that S-nitrosylation increases COX2 activity has been demonstrated using *in vitro* pathogen ^46^ and neurotoxicity ^47^ models, and in an *in vivo* model of myocardial infarction ^57^. Further studies demonstrate that iNOS inhibitors can block COX2 activity and its downstream pathogenic sequela, demonstrating a synergistic interaction between these two major inflammatory systems ^58,59^. Consistent with NOS2 and COX2 inflammatory pathway cross-talk in human breast cancer, a recent report finds co-expression of iNOS and COX2 predicts poor survival in breast cancer patients, and animal modeling confirms survival benefit with dual targeting of iNOS and COX2 ^58^. Thus, our work identifying antibodies that distinguish COX2 based on nitrosylation state in humans highlights the need for future investigations into the role that COX2 S-nitrosylation plays in breast cancer risk, progression and outcomes.

Main strengths of our study include the use of multiple, independent methods to investigate COX2 specificity of four different αCOX2 antibody clones; the use of robust biochemical approaches to demonstrate dependency of S-nitrosylation state on COX2 antibody recognition; and the inclusion of ~2000 breast cancer cases to assess dual COX2 staining. Of potential relevance to early onset breast cancer, we observed higher overall tumor cell COX2 staining and higher levels of S-nitrosylated COX2 in younger, primarily pre-menopausal age patients compared to that observed in the mostly postmenopausal patients enrolled in NHS1. This observation is consistent with previous reports demonstrating prostaglandin production and COX2 are positively regulated in mammary epithelial cells by ovarian hormones ^38,60^. One limitation of our study is the lack of inclusion of true normal breast tissue for assessing baseline levels of S-nitrosylated and non-nitrosylated COX2. Further, our assessment of COX2 levels in adjacent normal, DCIS and invasive cancers is based on a limited number of cases, requiring additional assessment.

To conclude, we find that commonly utilized antibodies directed against COX2 can distinguish between S-nitrosylated and non-nitrosylated forms of COX2. Further, we find that S-nitrosylated and non-nitrosylated COX2 have distinct subcellular distributions in both adjacent normal and breast and colon cancer tissues, providing evidence for distinct synthetic, trafficking, and functional pathways. As a result, previous work relying on COX2 IHC to define associations between COX2 expression and cancer parameters should be reviewed in light of these findings. Likewise, future COX2 studies should be designed with multiple antibody clones to detect both S-nitrosylated and non-nitrosylated forms of COX2. How both forms of COX2 are regulated, which tissue compartments express COX2 (e.g., epithelial, endothelial, immune), what subcellular locations they occupy, and how subcellular localization impacts COX2 function remain important, unanswered questions.

## Materials and Methods

### Ethics

Formalin-fixed paraffin-embedded (FFPE) human breast and colon tissue for this study was approved by the BWH/Harvard Cohorts Biorepository and Institutional Review Boards at Colorado Multiple Institution Review Board (COMIRB), and Oregon Health and Science University (OHSU). Written informed consent was given by participants when required.

### Human tissues

De-identified FFPE cases of breast (n=52) and colon (n=53) cancer tissue microarrays (TMAs) and breast cancer cases (n=2019) from the Nurses’ Health Study-1 (NHS1) ^61^ were obtained from the Channing Laboratory, Brigham and Women’s Hospital, Massachusetts. Young women’s FFPE breast cancer cases were acquired from the Young Women’s Breast Cancer Translational Program (YWBCTP) at the University of Colorado (n = 233). Breast tissue sections with adjacent normal, ductal carcinoma in situ (DCIS) and invasive ductal carcinoma on a single slide were obtained from Kaiser Permanente Northwest (KPNW) (n=10). A total of 233 YWBCTP and 1770 NHS1 cases were evaluated for dual COX2 IHC stain after exclusion of one entire TMA slide (249 cases) from the NHS1 cohort. The control cores for this TMA slide displayed staining several standard deviations above the average for the study, resulting in exclusion from analysis.

### Antigenic regions used for antibody generation

COX2 protein sequence for human (P35354), rat (P35355), and mouse (Q05769) were aligned with Uniprot (http://www.uniprot.org/). Antigenic peptide sequences used to generate each COX2 clone were obtained from the respective manufacturers and examined for differences in species and post-translational modification sites.

### Immunoblotting

Recombinant human COX1 protein (Abcam, Ab-198643, 4ng), human COX2 protein (Cayman Chemical, 60122, 4ng), and 25μg cell line lysates in RIPA buffer were separated by WES automated gel electrophoresis System (Protein Simple, San Jose, CA). Cell lines were procured from authenticated sources: mouse BRAFV600E melanoma (Wt) and COX1/2 CRISPR targeted sub-line^29^; human HCA-7 colon cancer (Sigma Aldrich #02091238), and human HCT-15 colon cancer (ATCC, # CCL-225). Primary antibodies and working concentrations were: COX2 SP21 clone (Thermo Fisher Scientific # RM-9121, at 25ng/uL), COX2 CX229 clone (Cayman Chemical # 160112, at 25ng/uL), COX2 CX294 clone (Agilent Dako # M3617, at 25ng/uL) and COX2 D5H5 clone (Cell Signaling Technology # 12282, at 25ng/uL), COX-1 (Cell Signaling Technology # 4841, at 25ng/uL) and GAPDH (14C10 clone, Cell Signaling Technology #2118, at 2ng/uL). For HRP-conjugated secondary antibodies and detection; anti-rabbit (Protein Simple # 042-206, RTU) or anti-mouse (Protein Simple# 042-205, RTU) were utilized, followed by chemiluminescent substrate (Protein Simple # PS-CS01, Luminol-S, Peroxide). Protein separation and signal detection utilized the WES system (Protein Simple, San Jose, CA) and immunoblot and electrophoretograms were composed and analyzed by Compass Software (Protein Simple, San Jose, CA). All lanes shown together in western data derive from the same experiment and were processed in parallel.

### S-nitrosylation and chemical de-nitrosylation

S-nitrosylation of proteins was detected by western blot using an S-nitrosylation specific antibody (HY8E12 clone, Abcam # 94930, 1:20) or the Pierce S-Nitrosylation Western Blot detection kit (Thermo Fisher Scientific # 90105). De-nitrosylation of proteins was performed using either 330 mM (low) or 1M (high) sodium (Na) ascorbate in HENS buffer (Thermo Fisher Scientific # 90106) as previously described ^47^.

### Immunohistochemical staining of FFPE tissues

Four μm sections of FFPE tissue were stained for single or dual COX2 IHC, or dual COX2 immunofluorescence (IF). Detailed protocols for staining are outlined in **Supplemental Table 4**. COX2 antibody clones were SP21 (Thermo Fisher scientific # RM-9121), CX229 (Cayman # 160112), and CX294 (Agilent Dako # M3617). Secondary antibodies and chromogens were Envision+ HRP detection (Agilent # K4001, # K4003) followed by 3,3’-Diaminobenzidine (DAB) (Agilent # K3468), or alkaline phosphatase detection (Enzo Life Sciences # ACC110-0150) followed by Warp Red (Biocare # WR806) for IHC staining and Alexa Fluor antibodies (Invitrogen # A11029, #A21245) for IF staining. IHC and IF stained slides were scanned using Aperio ScanScope AT (Leica Biosystems, Vista, CA) and Apotome (Zeiss, Jena, Germany) microscopes, respectively. IHC signal data werer captured and quantified using Aperio ImageScope analysis software (Leica Biosystems, Vista, CA) ^38^. All data acquisition was performed by investigators who were blinded to study group.

### Hierarchical and K means Clustering of SP21 and CX229 expression

Percent area stained for SP21 and CX229 dual stained FFPE tissue from the NHS1 and YWBCTP cohorts were separately subjected to Hierarchical and K-means clustering and optimal cluster numbers were obtained. Hierarchical clustering was performed using R studio software. For K means clustering, the lowest expression group was identified as the distribution containing the negative stained group above which all values would be considered positive ^62,63^.

### Statistical analysis

Comparisons for far/near adjacent normal, DCIS, and invasive cancer were done on GraphPad Prism 8 software using the two tailed t-test, with significance at P value of <0.05. Comparisons of clinical characteristics of the NHS1 and YWBCTP cohorts was performed using chi-squared test on GraphPad Prism 8 software.

## Data Availability

For western analysis data the associated Compass (Protein Simple, San Jose, CA) data files with raw data are available upon request. Other data that support the findings of this study are available from the corresponding author upon reasonable request.

## Acknowledgments

We thank Hadley Holden, and Marcelia Brown for immunohistochemistry and technical assistance; Sam Sivagnanam for assistance with hierarchical clustering analyses; Kristin Muessig for assistance with human subject research regulations, Christopher Rivard (University of Colorado, Denver) for COX2 knockout mouse tissue; Santiago Zelenay (CRUK Manchester Institute, Manchester UK) and Caetano Reis e Sousa (The Francis Crick Institute, London, UK) for the mouse melanoma cell line harboring a BRAFV600E mutation (Wt) and the sub-line deficient for COX1 and COX2; Naoki Oshimori for IF imaging; Lisa Coussens laboratory for insightful discussion on the project; and Weston Anderson for manuscript editing and development. We would like to thank the participants and staff of the YWBCTP at University of Colorado Hospital and NHS1 for their valuable contributions as well as the involved State Cancer Registries.

## Author Contributions

S.J, N.P, J.N, A.C and A.K performed IHC and cell culture experiment work, and P.S assisted with associated data analysis. M.R, R.M.T, and A.H.E performed analysis on the NHS-1 cohort. S.J, N.P, A.K, J.N, A.C, M.R, R.M.T, A.H.E, S.W, V.F.B and P.S contributed to discussion of the data. N.P, R.M.T, A.H.E, S.W and V.F.B read the manuscript and provided feedback. S.J and P.S led the project, interpreted the data, and wrote the manuscript. All authors approve the submitted version of the manuscript.

## Materials and Correspondence

Materials requests and correspondence may be addressed the corresponding author.

---

**Supplemental Figure 1.**
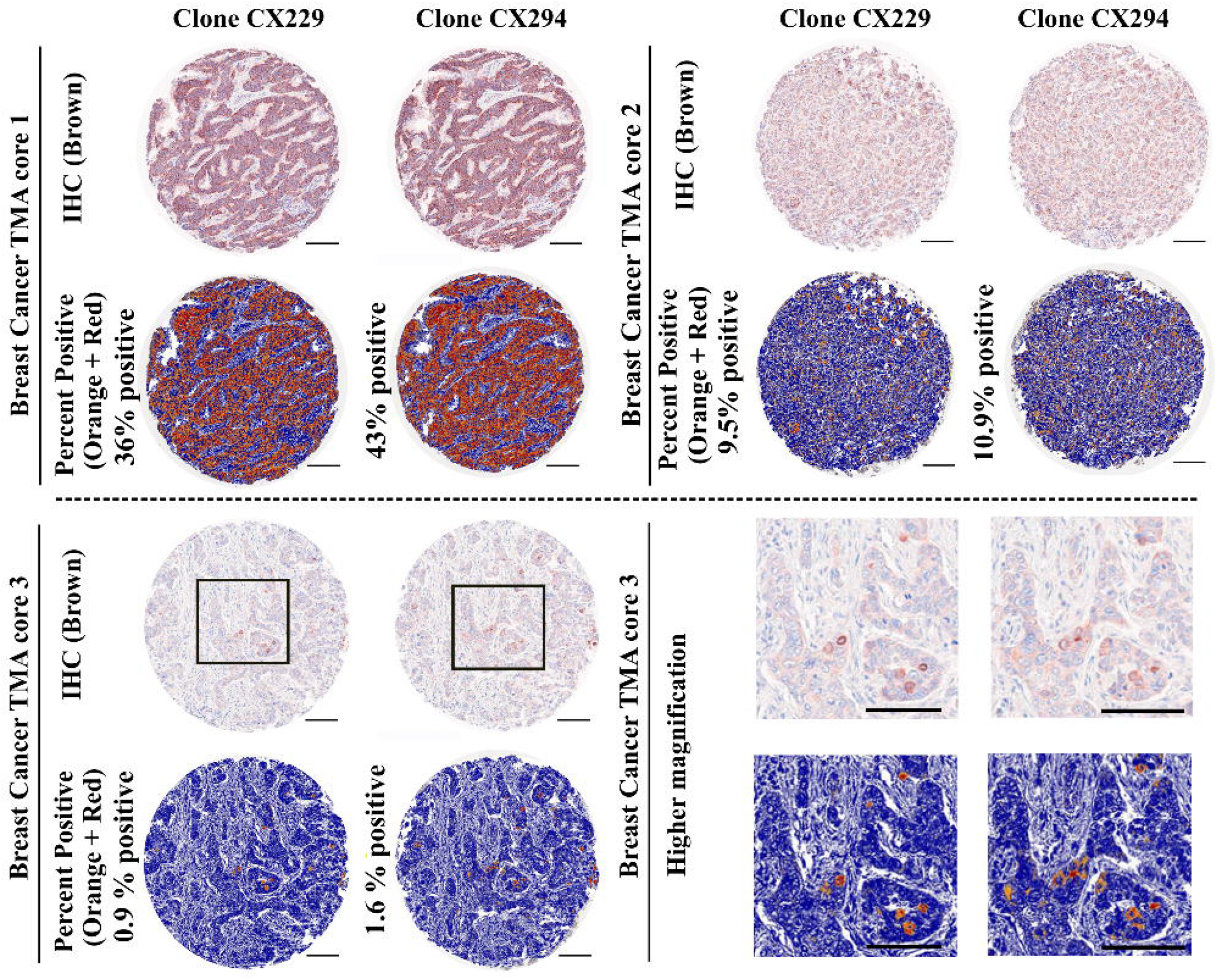
TMA cores from breast cancer cases stained for CX229 and CX294 have similar staining intensity, ranging from high (core #1), medium (core #2) or low (core #3). Higher magnification of core #3 (black box) shows that CX229 and CX294 have similar specificity in staining the same cells on sequential sections. Algorithmic analysis for each TMA core shows COX2 positive signal (orange and red) compared to negatively stained tissue (blue). Scale bar for all images is 100μm.

**Supplemental Figure 2.**
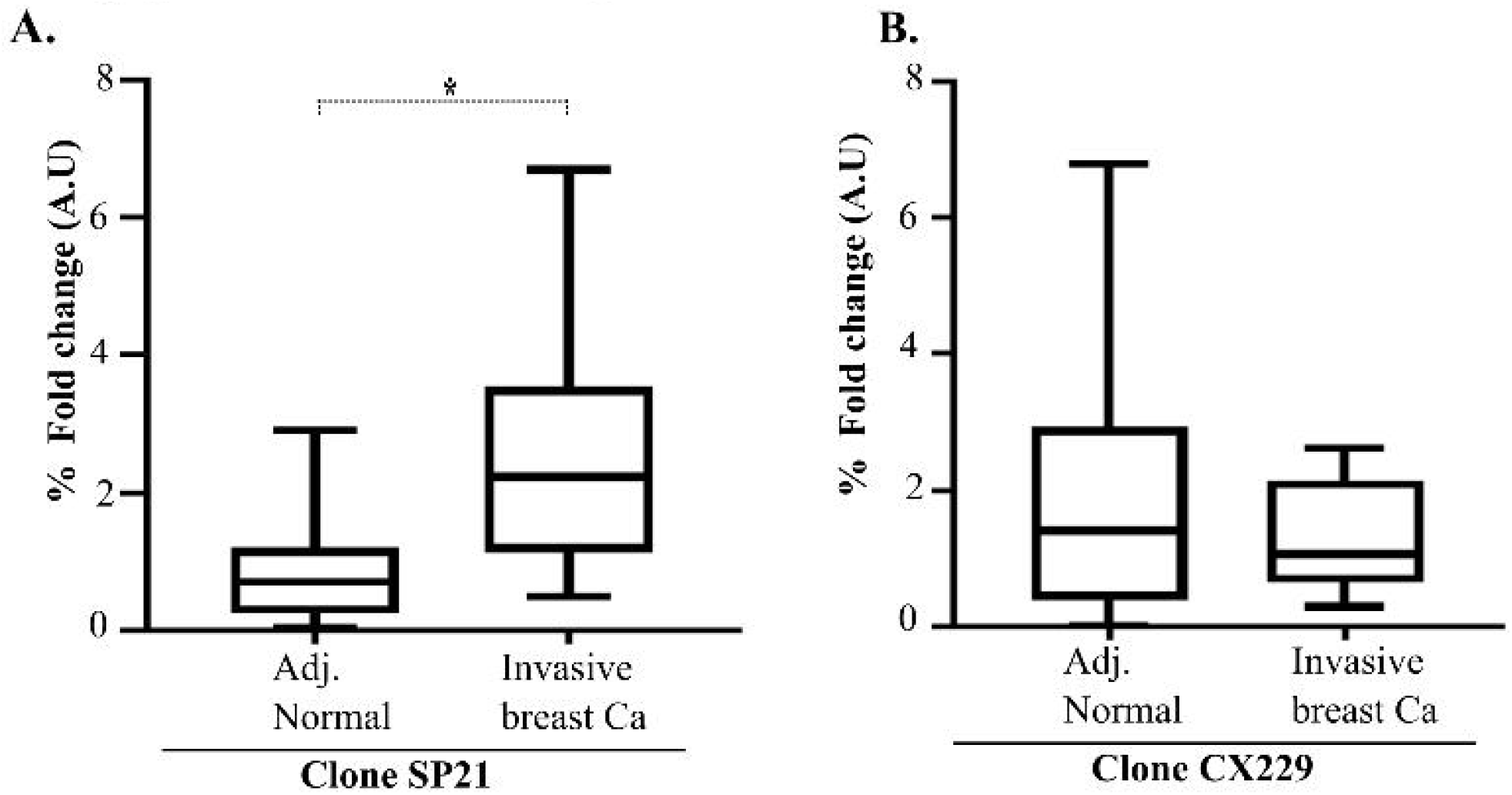
SP21 and CX229 clones show inverted staining patterns between adjacent normal and invasive cancer in Kaiser Pacific North West (KPNW) cases. **A**. SP21 staining intensity was low in adjacent normal breast tissue and increased in invasive breast cancer (blue circles, n=10, P value: *≤0.05). **B**. CX229 (blue triangles, n=10) shows highest COX2 expression in normal adjacent breast epithelium with a trend towards decreased COX2 expression in invasive breast cancer.

**Supplemental Figure 3.**
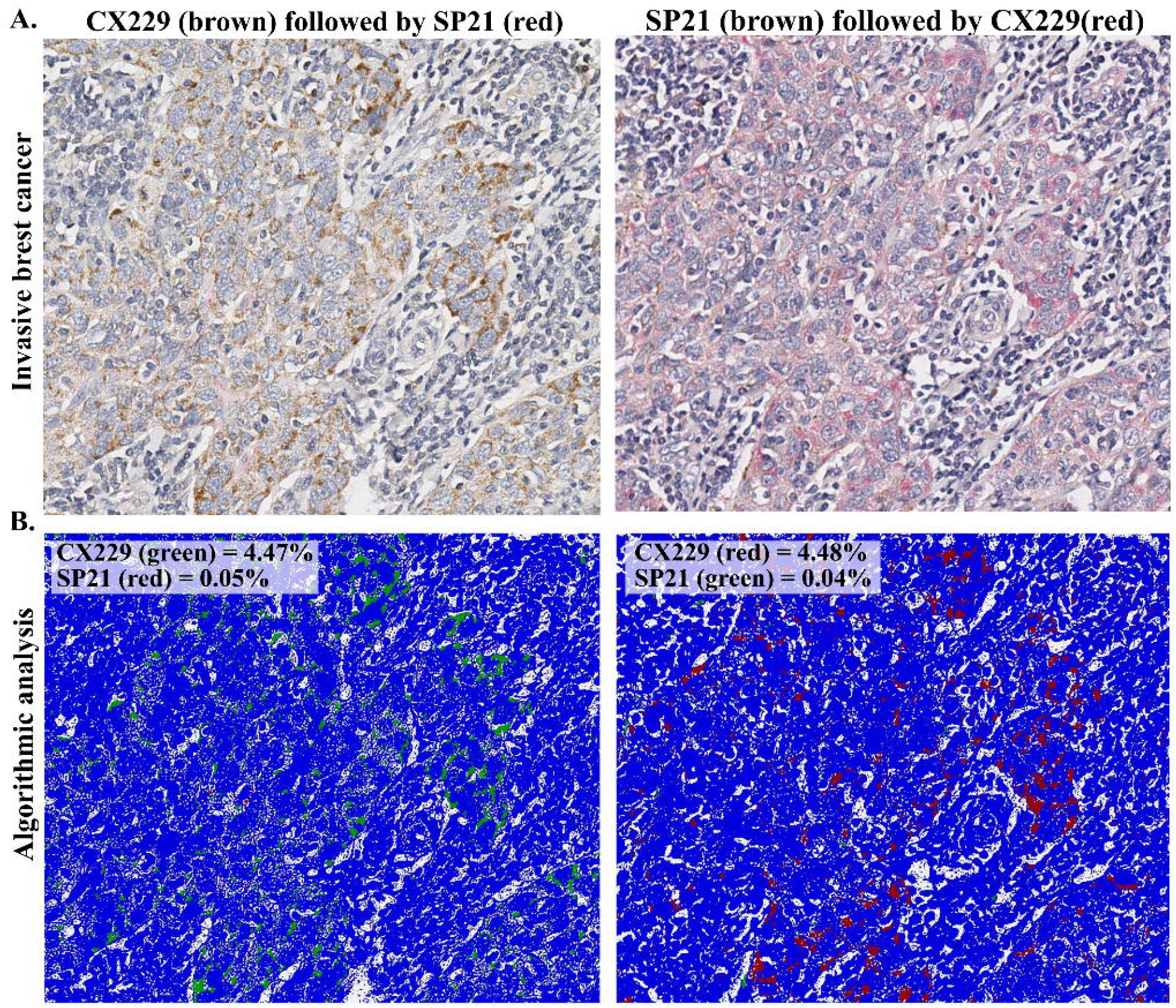
**A**. Dual IHC stained images of SP21 and CX229 show similar staining patterns when the order of antibodies and chromogens were switched during staining process. **B**. Algorithmic analysis of the images in part **A** of this figure (upper panel) revealed 4.47% CX229 (green) and 0.05% SP21 (red) expression (left, lower panel). With reversed antibody order, there was 4.48% CX229 (red) and 0.04% SP21 (green) expression (right, lower panel).

**Supplemental Figure 4.**
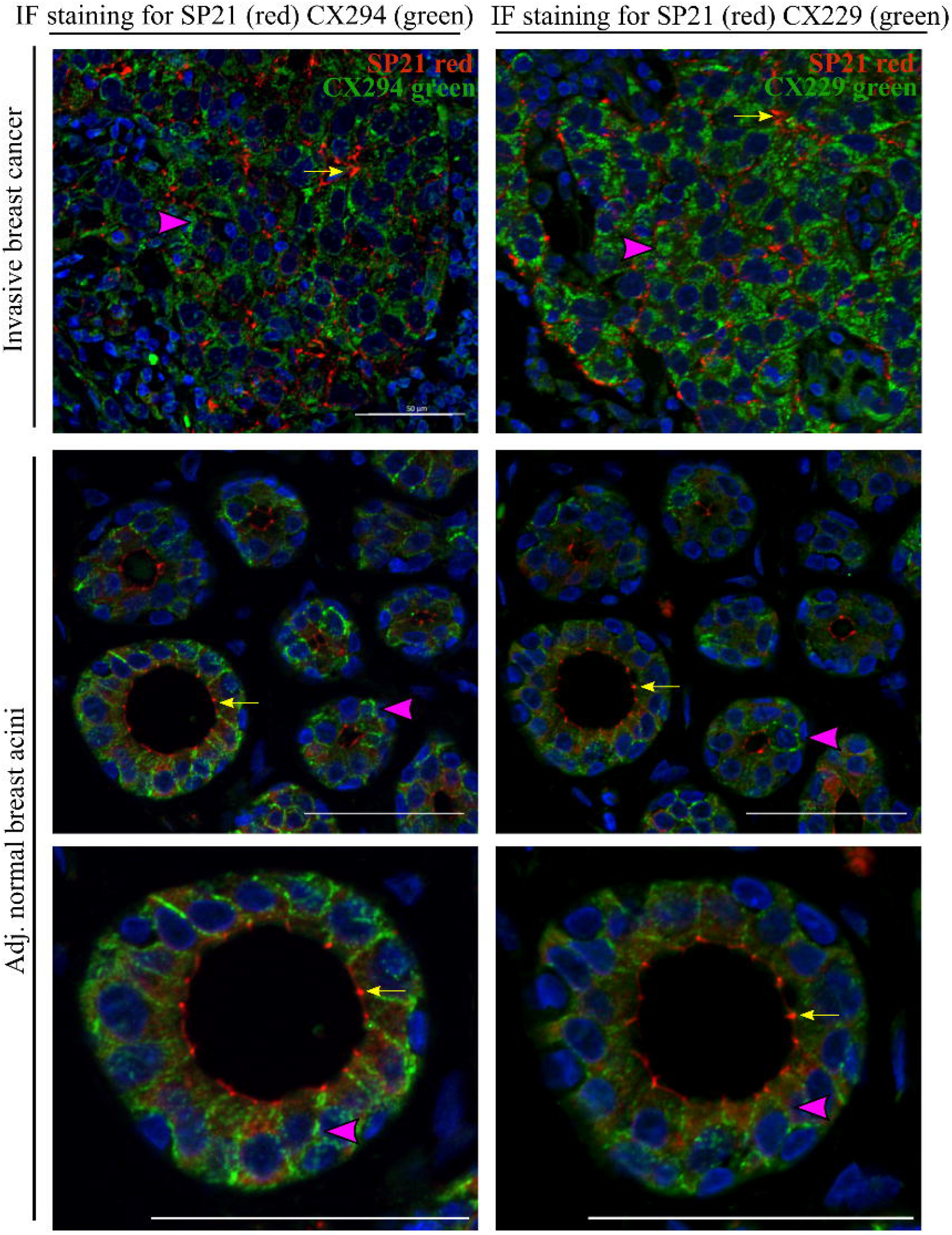
IF staining of serial sections of an invasive cancer case and adjacent normal breast acini show distinct sub-cellular localization of SP21, CX294 and CX229. The left column shows dual stains of SP21 and CX294 with minimal co-localization (SP21 = red, CX294 = green, co-localization = yellow). The right column shows dual stains of SP21 and CX229 with minimal co-localization (SP21 = red, CX294 = green, co-localization = yellow). Pink arrow heads show intense localized staining for CX294/CX229. Yellow arrows show intense localized staining for SP21. Scale bar for all images is 50μm.

