## Supplemental Tables for "S-nitrosylated and non-nitrosylated COX2 has differential expression and distinct subcellular localization in normal and breast cancer tissue"

| **TMA Staining Results** | **COX-2 Clones** | **COX-2 Positive Breast TMA (n=56)** |
| --- | --- | --- |
| Dual Positive COX-2 | CX229+/CX294+ | 53 (95%) |
| Positive for CX229 only | CX229+/CX294- | 0 |
| Positive for CX294 only | CX229-/ CX294+ | 0 |
| Dual negative COX-2 | CX229-/ CX294 - | 3 (5%) |

**Supplemental Table 1**: COX-2 expression in Breast TMA comparing COX2 clones CX229 and CX294

**Supplemental Table 2**: COX-2 expression in Breast and Colon TMA comparing clones SP21 and CX229

| **TMA Staining Results** | **COX-2 Clones** | **COX-2 Positive Breast TMA (n=52)** | **COX-2 Positive Colon TMA (n=53)** |
| --- | --- | --- | --- |
| Non-nitrosylated COX-2 only | CX229**+/**SP21**-** | 18 (35%) | 23 (43%) |
| Dual positive COX-2 | CX229**+/**SP21**+** | 29 (55%) | 21 (40%) |
| S-nitrosylated COX-2 only | CX229**-/**SP21**+** | 1 (2%) | 0 (0%) |
| Dual negative COX-2 | CX229**-/**SP21**-** | 4 (8%) | 9 (17%) |

**Supplemental Table 3.** Clinical characteristics of NHS 1 and YWBC cohorts

| **Clinical characteristics** | **Nurses’ Health Study 1** | **Young Women Breast Cancer** | **P value**  **(chi square test)** |
| --- | --- | --- | --- |
| **Overall N** | 1770 cases | 233 cases |  |
| **Age at dx, mean** | 57.8 | 38.24 |  |
| **Stage** |  |  |  |
| 1 | 885 (50%) | 67 (28.7%) |  |
| 2 | 600 (33.9%) | 97 (41.6%) | <0.0001 |
| 3 | 250 (14.1%) | 52 (22.3%) |  |
| 4 | 35 (2%) | 15 (6.52%) |  |
| **Grade** |  |  |  |
| 1 | 303 (17.1%) | 22 (9.4%) |  |
| 2 | 935 (52.8%) | 86 (36.9%) | <0.0001 |
| 3 | 416 (23.5%) | 109 (46.8%) |  |
| Missing | 116 (6.6%) | 16 (6.9%) |  |
| **Tumor size (cm)** |  |  |  |
| ≤1 | 439 (24.6%) | 30 (12.9%) |  |
| 1.1-2.0 | 682 (38.3%) | 73 (31.4%) | <0.0001 |
| 2.1-4.0 | 481 (27.1%) | 65 (27.9%) |  |
| >4.0 | 164 (9.8%) | 53 (22.7%) |  |
| Missing | 4 (0.2%) | 12 (5.1%) |  |
| **ER status** |  |  |  |
| Positive | 1361 (76.9%) | 142 (60.9%) |  |
| Negative | 404 (22.8%) | 87 (37.3%) | <0.0001 |
| Missing | 5 (0.3%) | 4 (1.8%) |  |
| **PR status** |  |  |  |
| Positive | 1137 (64.2%) | 135 (57.9%) |  |
| Negative | 625 (35.3%) | 93 (39.9%) | 0.1156 |
| Missing | 8 (0.5%) | 5 (2.1%) |  |
| **HER2 status** |  |  |  |
| Positive | 195 (11.0%) | 67 (28.7%) | <0.0001 |
| Negative | 1546 (87.4%) | 147 (63%) |  |
| Missing | 29 (1.6%) | 19 (8.3%) |  |
| **Hormone therapy** |  |  |  |
| Yes | 837 (47.4%) | 79 (33.9%) |  |
| No | 433 (24.6%) | 67 (28.8%) | 0.0047 |
| Missing | 496 (28%) | 87 (37.3%) |  |
| **Radiation** |  |  |  |
| Yes | 545 (30.7%) | 90 (38.6%) |  |
| No | 735 (41.5%) | 55 (43.5%) | <0.0001 |
| Missing | 490 (27.8%) | 88 (37.8%) |  |
| **Chemotherapy** |  |  |  |
| Yes | 492 (27.8%) | 137 (58.9%) |  |
| No | 772 (43.6%) | 19 (8.1%) | <0.0001 |
| Missing | 506 (28.6%) | 77 (33%) |  |

**Supplemental Table4**: The methodological details for IHC and IF staining

| **Stain/single or dual** | **Antigen retrieval** | **Peroxidase and Protein Blocks** | **Primary Antibody** | **Secondary Antibody** | **Chromogen** |
| --- | --- | --- | --- | --- | --- |
| **IHC –**  **SP21 stain** | 125⁰C under pressure for 5 mins with TRS (Dako  #S1699) | Peroxidase block (1ml 3% H2O2 + 9 ml methanol) for 10 mins   Protein block (Biocare  # BS966L) for 10 mins | SP21 clone (Thermoscientific # RM-9121) 1:200 for 60 mins | Envision + HRP antibody (Dako # K4003 rabbit) for 30 mins | DAB (Dako # K3468) for 10 mins |
| **IHC- CX229**  **stain** | 125⁰C under pressure for 5 mins with TRS (Dako  #S1699) | Peroxidase block (1ml 3% H2O2 + 9 ml methanol) for 10 mins   Protein block (Biocare  # BS966L) for 10 mins | CX229 clone (Cayman, # 160112) 1:250  for 60 mins | Envision + HRP antibody (Dako # K4001 mouse) for 30 mins | DAB (Dako # K3468) for 10 mins |
| **IHC- CX294**  **stain** | 125⁰C under pressure for 5 mins with TRS (Dako  #S1699) | Peroxidase block (1ml 3% H2O2 + 9 ml methanol) for 10 mins   Protein block (Biocare  # BS966L) for 10 mins | CX294 clone (Agilent # M3617) 1:50  for 60 mins | Envision + HRP antibody (Dako # K4001 mouse) for 30 mins | DAB (Dako # K3468) for 10 mins |
| **IHC- Dual SP21 and CX229**  **stain** | 125⁰C under pressure for 5 mins with TRS (Dako  #S1699) | *1st antibody stain*: Peroxidase  (1ml 3% H2O2 + 9 ml  methanol) for 10 mins | CX229 clone (Cayman, #160112) 1:250for 60 mins | Envision + HRP antibody (Dako # K4001mouse) for 30 mins | DAB (Dako # K3468) for 10 mins |
|  |  | Protein block (Biocare  # BS966L) for 10 mins  *2nd antibody stain*: Protein block (Biocare # BS966L) for 10 mins |  |  |  |
